# Function and essentiality of *Plasmodium falciparum* plasmepsin V

**DOI:** 10.1101/404798

**Authors:** Nonlawat Boonyalai, Christine R. Collins, Fiona Hackett, Chrislaine Withers-Martinez, Michael J. Blackman

## Abstract

The malaria parasite replicates within erythrocytes. The pathogenesis of clinical malaria is in large part due to the capacity of the parasite to remodel its host cell. To do this, intraerythrocytic stages of *Plasmodium falciparum* export more than 300 proteins that dramatically alter the morphology of the infected erythrocyte as well as its mechanical and adhesive properties. *P. falciparum* plasmepsin V (PfPMV) is an aspartic protease that processes proteins for export into the host erythrocyte and is thought to play a key role in parasite virulence and survival. However, although standard techniques for gene disruption as well as conditional protein knockdown have been previously attempted with the *pfpmv* gene, complete gene removal or knockdown was not achieved so direct genetic proof that PMV is an essential protein has not yet been established. Here we have used a conditional gene excision approach combining CRISPR-Cas9 gene editing and DiCre-mediated recombination to functionally inactivate the *pfpmv* gene. The resulting mutant parasites displayed a severe growth defect. Detailed phenotypic analysis showed that development of the mutant parasites was arrested at the ring-to-trophozoite transition in the erythrocytic cycle following gene excision, likely due to a defect in protein export. Our findings are the first to elucidate the effects of PMV gene disruption, showing that it is essential for parasite viability in asexual blood stages. The mutant parasites can now be used as a platform to further dissect the *Plasmodium* protein export pathway.

## Introduction

Malaria was responsible for approximately 445,000 deaths and 216 million clinical cases in 2016, an increase of ∼5 million cases over the previous year [1]. Vital to the growth and pathogenicity of the parasite is host cell remodelling in which the parasite modifies the host erythrocyte by the synthesis and export of over 300 parasite proteins beyond the bounds of the parasitophorous vacuole (PV) within which it replicates (for reviews, see [2,3]). The exported proteins are trafficked from the parasite to the host erythrocyte via a putative parasite-derived protein complex known as the translocon of exported proteins (PTEX), located within the PV membrane (PVM) [4,5]. The exported proteins extensively alter the mechanical and adhesive properties of infected erythrocytes, resulting in vascular sequestration of the infected cells and eventual destruction of the host erythrocyte [6]. Protein export in *Plasmodium* has been most intensively studied in *P. falciparum* (reviewed in [7,8]), where the majority of known exported proteins contain the pentameric localisation motif RxLxE/Q/D, termed *Plasmodium* export element (PEXEL), typically located downstream of the N-terminal secretory signal sequence that regulates entry into the ER [9,10]. Some proteins lacking a PEXEL motif, referred to as PEXEL-negative exported proteins (PNEPs), can also be exported [11]. PNEPs do not contain a typical secretory signal sequence so are unlikely to be N-terminally processed, but the first 20 amino acid residues of PNEPs are sufficient to mediate their export. The unfolding of soluble PNEPs is then required for trafficking into the host cell [12].

*P. falciparum* plasmepsin V (PfPMV) is an ER-located aspartic protease comprising 590 amino acid residues (∼68 kDa) [13]. PfPMV expression levels in the parasite progressively increase throughout schizogony [13]. Numerous studies have now established that PMV is directly responsible for cleavage of the PEXEL motif within exported proteins. Cleavage occurs on the C-terminal side of the conserved Leu residue (RxL↓), revealing a new N-terminus that is rapidly acetylated (^Ac^-xE/Q/D) [14]. A recent x-ray crystal structure of *Plasmodium vivax* PMV (PvPMV) revealed a canonical aspartyl protease fold with several important features [15]. These include a nepenthesin (NAP)-like insertion within the N-terminal part of the enzyme, which may control substrate entry into the active site and influence enzyme specificity, as well as a helix-turn-helix (HTH) motif near the C-terminus of the enzyme, which is conserved in PMV from other *Plasmodium* species but not found in other parasite plasmepsins involved in haemoglobin digestion. PMV also possesses an unpaired Cys residue (C140 in PvPMV, equivalent to C178 in the *P. falciparum* enzyme) which is located in the flap of the structure and is restricted to *Plasmodium* species. In attempts to establish the druggability and function of PMV, several peptidomimetic inhibitors based on the PEXEL motif have been developed [16-18]. A co-crystal structure of PMV bound to one of these compounds, the PEXEL-mimetic WEHI-842, showed that although the unpaired Cys points into the active site of the enzyme, it does not appear to make contact with the inhibitor [15,19]. Another inhibitor, WEHI-916, inhibits the activity of purified PMV isolated from both *P. falciparum* and *P. vivax*, and higher concentrations of WEHI-916 were required to kill parasites engineered to over-express PfPMV indicating on-target efficacy [16,18]. Compound 1, a hydroxyl-ethylamine PEXEL-mimetic, inhibited PfPMV activity in vitro with picomolar potency but failed to block parasite growth due to poor stability and membrane permeability [17]. Collectively, these findings indicate that PMV has a number of distinguishing features that could be exploited in drug design, and can potentially be targeted with suitable inhibitory compounds.

An important component in the validation of any enzyme as a drug target is genetic ablation of enzyme expression. Unfortunately, previous attempts to genetically delete or knock-down PMV expression have been only partially successful. Work by Boddey et al [20] failed to disrupt the *P. berghei* PMV (PbPMV), suggesting (but not proving) that the gene is essential. Russo et al [21] similarly attempted to disrupt the *pfpmv* gene using an allelic replacement approach. Single crossover homologous recombination into the endogenous PMV locus was possible only when the catalytic dyad aspartate codon was preserved, whilst four separate transfection experiments with constructs designed for non-synonymous alteration of the active site codon were not successful. Conditional knockdown of PfPMV expression was also attempted using the RNA-degrading *glmS* ribozyme system [18]. Although this resulted in around 75-90% knockdown of PMV expression, the remaining PMV levels were presumably sufficient to enable export and sustain parasite development. Gambini et al [17] generated an inducible PMV knock-down by fusing PMV with a destabilising domain (DD), but only a 4 to 10-fold knock-down of PMV cellular levels was achieved using this approach, which did not affect parasite viability. In summary, whilst some of these data are consistent with an essential role for PMV, in the absence of any reports of complete ablation of PfPMV expression, direct genetic proof that PMV is an essential protein is lacking.

Here we have used a robust conditional genetic approach to truncate the *pfpmv* gene. Our findings are the first to confirm genetically that deletion of PfPMV has a significant effect on ring-to-trophozoite development and ultimately results in parasite death.

## Results

### Generation of modified *pfpmv* parasites using CRISPR-Cas9

Previous attempts to disrupt the *pfpmv* gene using conventional genetic techniques were unsuccessful [20,21], and conditional knockdown approaches did not significantly affect PEXEL processing or parasite viability [17,18], presumably due to relatively low levels of PfPMV expression being sufficient to sustain parasite viability. To explore the consequences of complete functional inactivation of PMV, we therefore took advantage of the DiCre conditional recombinase system, recently adapted to *P. falciparum* [22]. Using Cas9-mediated genome editing [23] we first introduced synthetic introns containing *loxP* sites [24] into the endogenous *pfpmv* locus such that they flanked (floxed) an internal segment of the gene encoding Asp133 to Thr590. At the same time, the modified gene was fused to a C-terminal HA3 epitope tag, as well as a 2A sequence followed by the aminoglycoside 3′-phosphotransferase (*neo-R*) gene sequence conferring neomycin resistance activity to enable selection for integration events (Fig 1A). Importantly, one of the PfPMV catalytic dyad residues (Asp365) is included within the floxed region. The repair plasmid pT2A-5´UTR-3´-PMV-ΔDHFR was based on pT2A-DDI-1cKO, which contains a modified selection-linked integration (SLI) region [25]. The genomic modification was made in the DiCre-expressing *P. falciparum* B11 parasite clone [26] such that excision of the floxed sequence could be induced by treatment of the transgenic parasites with rapamycin (RAP). DiCre-mediated excision was predicted to generate an internally-truncated mutant form of PMV lacking one of the catalytic dyad residues. Excision would also remove the *neo-R* gene and induce expression of GFP as a fluorescence reporter indicative of parasites expressing the truncated *pfpmv* gene (Fig 2A).

**Fig 1.**
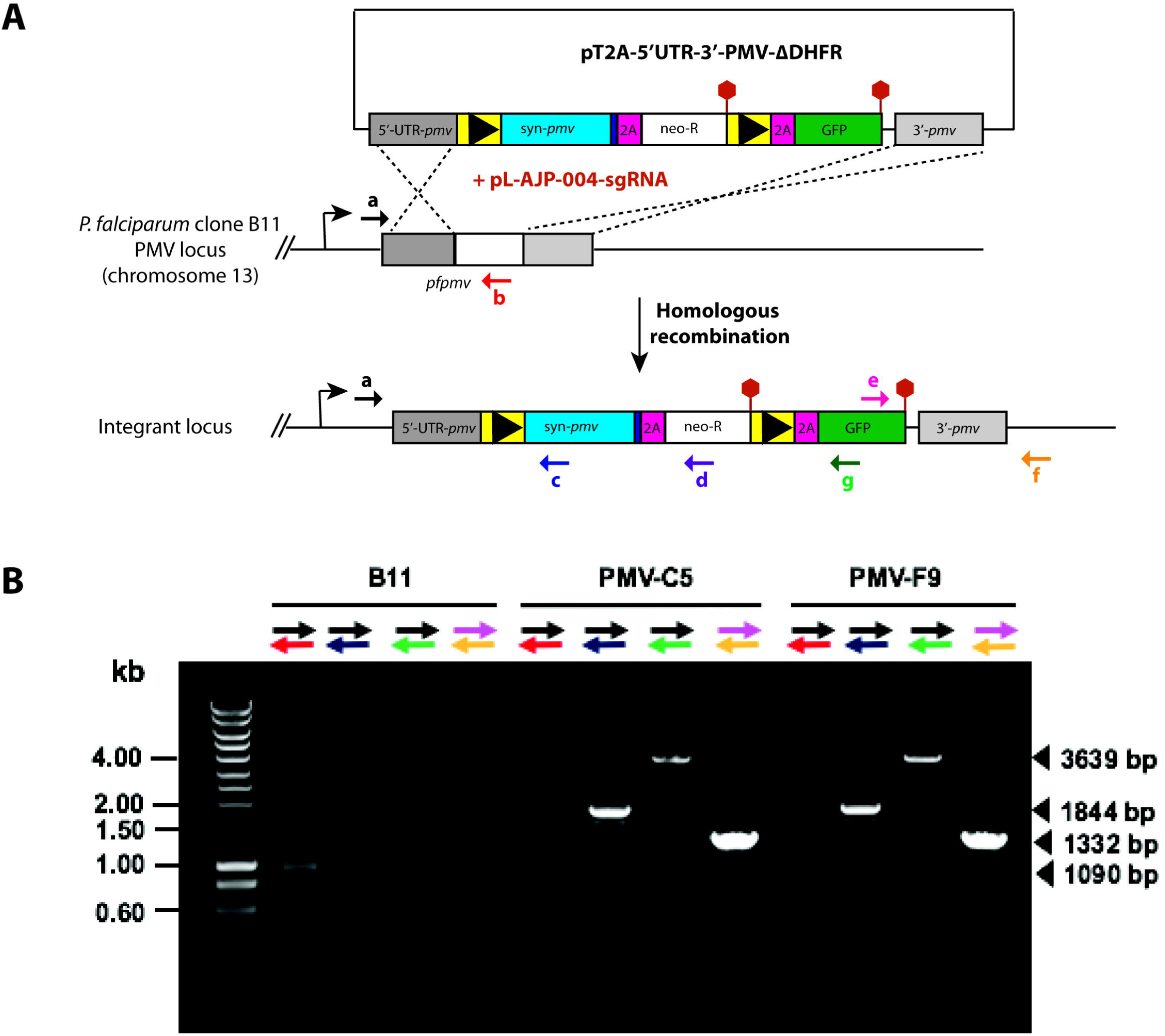
Generation of *P. falciparum* parasite lines expressing PfPMV-HA. (A) Using Cas9-mediated recombination, the region of the *pfpmv* gene encoding Asp133 to Thr590 was replaced with two *loxP* (black arrowhead)-containing *P. falciparum* SERA2 introns (yellow box) flanking a recondonised *pfpmv* (blue box) fused to a *3xHA* epitope tag (purple box), a *2A* sequence (pink box), and a *neo-R* gene (white box) and stop codon (red hexagon). The second *loxPint* was also fused with *2A* and *gfp* gene sequences. The *gfp* gene is translated only following site-specific recombination of the *loxP* sites by DiCre recombinase. Positions of hybridisation of primers used for confirmation of the integration event by diagnostic PCR are shown as coloured arrows. (B) Diagnostic PCR analysis of genomic DNA of the control parental B11 and integrant *P. falciparum* clones, confirming the predicted homologous recombination event.

**Fig 2.**
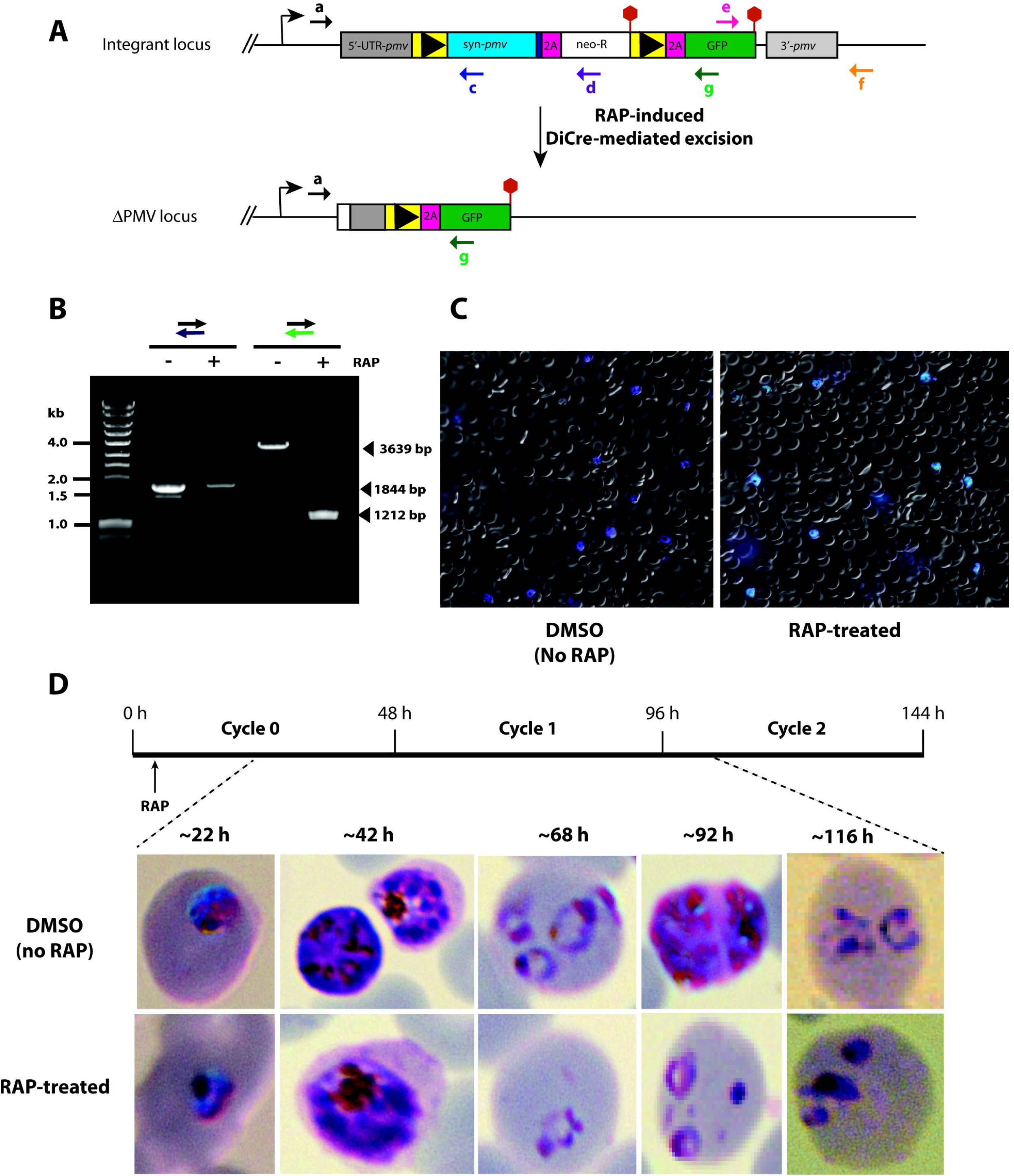
DiCre-mediated conditional disruption of *P. falciparum* PMV expression. (A) Strategy for conditional truncation of the *pfpmv* gene. Positions of hybridisation of primers used for diagnostic PCR analysis of the integration and excision events are shown as coloured arrows. (B) Diagnostic PCR analysis of genomic DNA from transgenic *P. falciparum* PMV-C5 line at 24 h in cycle 0 (∼20 h post RAP- or DMSO-treatment), confirming the predicted DiCre-mediated excision events. Expected sizes of the PCR amplicons specific for the intact or excised locus are indicated. (C) Live GFP fluorescence assay of cycle 0 PMV-C5 parasites ∼42 h following treatment at ring stage with DMSO or RAP. Parasite nuclei were stained with Hoechst 33342. (D) Giemsa-stained blood smears, showing the morphology of treated PMV-C5 parasites. The time-course of treatment and subsequent monitoring of the cultures is indicated (top).

Successful modification of the *pfpmv* gene in the transfected parasite population following the introduction of the targeting vector was confirmed by diagnostic PCR (data not shown). Limiting dilution cloning of the modified parasites resulted in the isolation of parasite clones PMV-C5 and PMV-F9, which were derived from independent transfections using different guide RNAs. Modification of the native *pfpmv* locus was confirmed in both transgenic parasite lines by diagnostic PCR (Fig 1B). Both clones replicated at a rate indistinguishable from that of the parental B11 parasites, suggesting that the modifications did not affect parasite viability. Transgenic parasite clone PMV-C5 was used for all subsequent experiments.

### Conditional truncation of the *pfpmv* gene leads to a developmental arrest

Expression of the recodonised *pfpmv* gene in transgenic parasite clone PMV-C5 was expected to produce an epitope-tagged PMV product (called PMV-HA), as well as expression of the *neo-R* gene product. Note that in this event GFP is not produced because of the presence of a translational stop codon directly downstream of the *neo-R* gene. Upon RAP-treatment, site-specific recombination between the introduced *loxP* sites in the modified *pfpmv* locus of the PMV-C5 parasites was anticipated to reconstitute a functional, albeit chimeric, intron. Splicing of this chimeric intron results in a truncated form of PMV lacking the HA epitope tag, as well as allowing expression of free GFP. DNA extracts of RAP-treated and mock-treated (DMSO-treated control) PMV-C5 parasites were analysed by diagnostic PCR (Fig 2A). To confirm whether RAP-induced DiCre-mediated excision took place as expected, we monitored the appearance of GFP-positive parasites following treatment of synchronous ring-stage parasites with DMSO or RAP. As expected, GFP fluorescence was only detected in the RAP-treated parasites (Fig 2C). Indeed, no parasites lacking GFP fluorescence could be detected upon microscopic examination of the RAP-treated cultures, indicating highly efficient excision of the floxed *pfpmv* sequence. This was confirmed by FACS analysis (Fig 3B), showing that 98.86 % of the RAP-treated parasites displayed GFP fluorescence. These results confirmed the PCR-derived excision data and demonstrated essentially complete conditional truncation of PMV within a single erythrocytic cycle in the PMV-C5 parasite clone.

**Fig 3.**
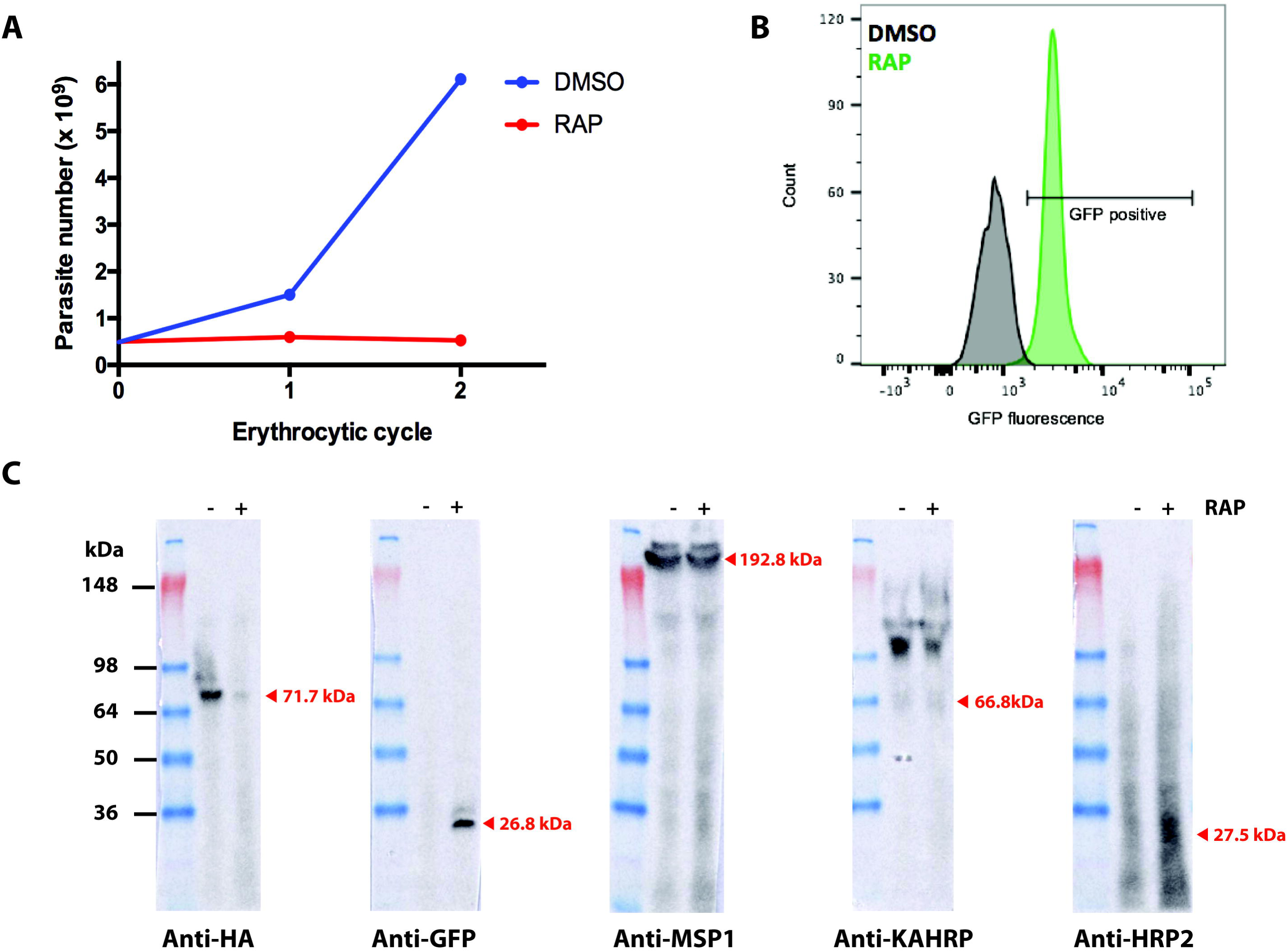
Functional disruption of PfPMV results in developmental arrest at the ring-to-trophozoite transition. (A) Growth assay showing relative replication rates over the two erythrocytic cycles following treatment of PMV-C5 ring stages with DMSO or RAP. Parasite number values were determined by FACS as described in Materials and Methods. (B) GFP fluorescence at ∼42 h (schizonts) of the PMV-C5 line treated at ring stage with DMSO (black) or RAP (green) as determined by flow cytometry of fixed cultures. The percentage of GFP-positive RAP-treated parasites (i.e. the excision rate) was 98.86%. (C) Western blot analysis of clone PMV-C5 following mock-treatment (-RAP) or RAP-treatment (+RAP). Schizont extracts (∼42 h post-treatment) were probed with antibodies to detect HA-tagged PfPMV (anti-HA), or antibodies to GFP, MSP1, KAHRP, or HRP2.

To initially explore the effects of PMV truncation on parasite viability, we examined growth and development of the ΔPMV mutant by microscopic examination of Giemsa-stained cultures. This revealed that although RAP-treated PMV-C5 parasites appeared morphologically normal throughout the erythrocytic growth cycle in which the parasites were treated (henceforward referred to as cycle 0), development of the mutant parasites stalled at ring stage in the following cycle (cycle 1; Fig 2D). By the end of cycle 1, only arrested pycnotic forms were observed. Collectively, these results suggested that ablation of PMV expression causes a severe growth defect and developmental arrest between ring and trophozoite stage.

### Egress, Invasion and Protein export are not affected by disruption of the *pfpmv* gene in cycle 0

Whilst the above results showed that truncation of PfPMV affected parasite development in cycle 1, it did not rule out the possibility of defects in egress and invasion of fresh erythrocytes at the end of cycle 0 (the cycle of RAP-treatment). To evaluate and quantify this, we used flow cytometry to monitor increases in parasitaemia at the transition between cycle 0 schizonts and cycle 1 rings. As shown in Fig 3A, the parasitaemia of PMV-C5 cultures at the end of cycle 0 as well as at the early ring stage of cycle 1 was unaffected by RAP-treatment, indicating no effects of PMV ablation on schizont development in cycle 0, or the egress and invasion capacity of the released merozoites. However, upon further development of the new parasite generation in cycle 1, the parasitaemia in the RAP-treated culture reached plateau after the ring-trophozoite transition. This confirmed that the truncation of PfPMV did not cause any effects on egress and invasion and that the enzyme is essential for the ring to trophozoite developmental transition of the intracellular parasite. Confirmation of the high excision rate of the PMV-C5 clone following RAP-treatment was also confirmed by FACS analysis (Fig 3B).

To further explore the PMV-null phenotype, we examined the expression of PMV-HA as well as other non-exported and exported parasite proteins. Extracts of DMSO and RAP-treated PMV-C5 parasites were analysed by immunoblot ∼42 h following treatment, using antibodies specific for the HA epitope tag as well as GFP, the merozoite plasma membrane surface protein MSP1, and the exported proteins KAHRP and HRP2. As shown in Fig 3C, only a faint HA signal was detected in the RAP-treated sample, whilst a corrrespondingly strong GFP signal was observed only in this sample. These data were in good agreement with the diagnostic PCR results, indicating high efficiency of conditional excision of the floxed sequence in the PMV-C5 clone. In contrast, no differences between DMSO- and RAP-treated parasites were detectable in expression levels of MSP1 (thought to play an important role in egress and invasion [27]) Similarly, there were no significant differences in expression of KAHRP and HRP2, both of which are examples of PEXEL-containing proteins that are cleaved by PMV [20,21,28]. This is consistent with the notion that protein export remains unaffected in cycle 0 in the RAP-treated parasites.

We next determined the effects of PfPMV truncation on its subcellular localisation within the parasite, as well as on the trafficking of other exported proteins in cycle 0. Immunofluorescence analysis (IFA) showed that, as expected, PMV-HA showed a perinuclear localisation in schizonts of control PMV-C5 parasites but the signal was lost in RAP-treated parasites (Fig 4). To determine the effects of PMV ablation on protein export, mock- and RAP-treated PMV-C5 parasites were probed with anti-KAHRP2 and anti-HRP2 antibodies. The localisation and intensity of the MSP1 signal and the KHARP signal did not alter upon RAP-treatment, correlating well with the western blot analysis. The HRP2 signal remained detectable in both control and the RAP-treated parasites, but in contrast to the situation with the other proteins examined, the fluorescence intensity of the HRP2 signal was substantially decreased in RAP-treated parasites, suggesting a partial effect of PMV depletion on HRP2 trafficking. Together, these data confirmed the loss of PMV-HA in RAP-treated parasites whilst indicating that both exported proteins (KAHRP and HRP2) and merozoite surface proteins still localise correctly in cycle 0. However, the decreased HRP2 signal intensity was consistent with a partial impact on protein export in cycle 0.

**Fig 4.**
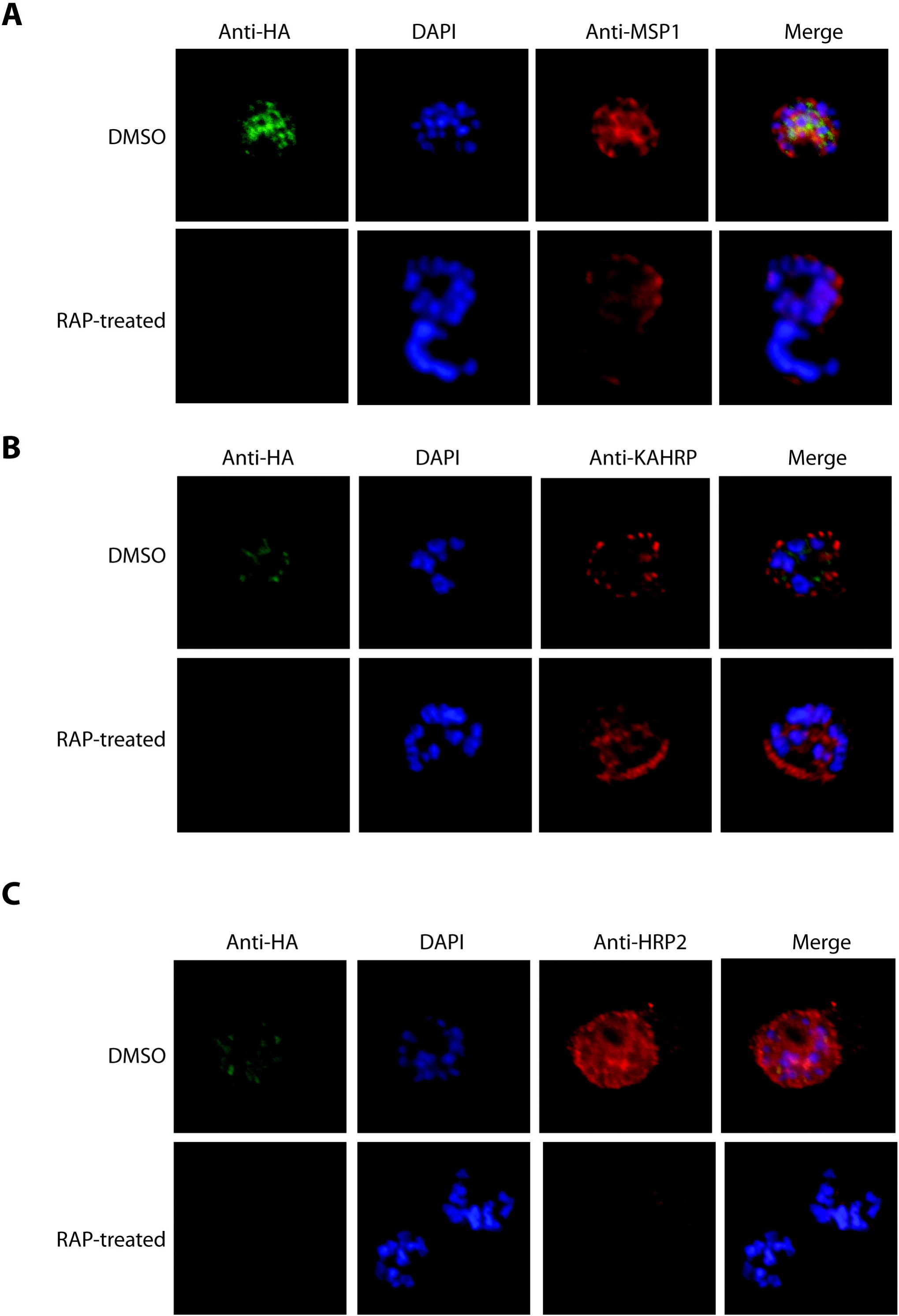
IFA of schizonts of control (DMSO-treated) and RAP-treated integrant clone PMV-C5 ∼42 h following treatment (cycle 0). The PMV-HA signal was lost following RAP treatment. Localization of MSP1, KAHRP and HRP2 was unaffected; however, the HRP2 signal was significantly decreased in intensity. Parasite nuclei were visualised by staining with DAPI.

## Discussion

Malarial proteolytic enzymes play regulatory and effector roles in multiple key biological processes in this important pathogen and have long been of interest as potential drug targets. In this study, we have shown for the first time that functional ablation of the *pfpmv* gene leads to a block at the ring-to-trophozoite transition and death of the parasite. To achieve this, we employed the robust conditional DiCre approach in combination with Cas9-induced gene modification. This system provides a rapid and efficient means of generating transgenic parasites as well as enabling complete disruption of the gene of interest [29,30]. Our study demonstrates that DiCre-mediated conditional knockout is a powerful tool to study essential genes in the human malaria parasite. Our genetic strategy also incorporated a method to permit the selection of parasites in which genomic integration of the input constructs had taken place, termed selection-linked integration (SLI), previously developed by Birnbaum *et al* [25]. For this, our rescue plasmid contained an SLI-resistance marker (neomycin phosphotransferase), which was linked to the modified *pfpmv* gene separated by a 2A ‘skip’ peptide [31,32]. This approach allows selection for correct integration by one to two weeks of neomycin selection. In our case, with the assistance of Cas9-mediated integration we in fact did not need to use neomycin selection in our experiments to select for correct integration. However, our plasmids could be useful for future genetic complementation studies.

When the PMV-modified parasite clone PMV-C5 was RAP-treated at early ring stage for just 3 h, followed by removal of RAP, parasite development proceeded normally in the first erythrocytic cycle (cycle 0). Whilst expression of most of the parasite proteins examined appeared unaffected, we did observe some impact on expression of the exported parasite protein HRP2. This is reminiscent of the observation by Russo et al [21] that the levels of HRP2 and RESA (ring-infected erythrocyte surface antigen) were reduced by 30-50% in PMV mutants. We suspect that this may be due to low levels of PMV expression early in cycle 0, perhaps due to transcription early in the erythrocytic cycle prior to gene excision. That low levels of PMV are sufficient to sustain development has also been indicated by previous conditional protein knockdown experiments [18]. Egress and invasion at the end of cycle 0 was not affected in the RAP-treated PMV-C5 parasites; however, parasite development in the next cycle stalled at the ring-to-trophozoite transition. This phenotype is very similar to that observed upon treatment with the PMV inhibitor WEHI-916 [18], where the drug-treated parasites showed a growth defect at the ring-to-trophozoite transition from approximately 20 h post-invasion. A similar parasite developmental arrest at ring stage was also observed following conditional ablation of the PTEX component HSP101 [33,34]. This implies that PMV function may be associated with PTEX, and may be essential for ring-to-trophozoite development.

In summary, using a DiCre-mediated conditional gene editing approach to selectively disrupt the *pfpmv* gene, we have shown that the gene is essential for the ring-to-trophozoite transition of intracellular growth. This engineered platform will be useful for further study of PEXEL-protein export as well as for dissection of PMV domain interactions, providing further impetus for focusing on PMV as a new potential antimalarial drug target.

## Materials and Methods

### *P. falciparum* culture, transfection and limiting dilution cloning

Parasites (wild type clone 3D7 and the DiCre-expressing clone B11 [26]) were routinely cultured at 37ºC in human erythrocytes at 1-4% haematocrit in RPMI 1640 (Life Technologies) supplemented with 2.3 gL^−1^ sodium bicarbonate, 4 gL^−1^ dextrose, 5.957 gL^−1^ HEPES, 0.05 gL^−1^ hypoxanthine, 0.5% (w/v) Albumax II, 0.025 gL^−1^ gentamycin sulphate, and 0.292 gL^−1^ L-glutamine (complete medium) in an atmosphere of 90% nitrogen, 5% carbon dioxide and 5% oxygen [35,36]. Routine microscopic examination of parasite growth was performed by fixing air-dried thin blood films with 100% methanol before staining with 10% Giemsa stain (VWR international) in 6.7 mM phosphate buffer, pH 7.1. For synchronization, mature schizont stage parasites were isolated on cushions of 70% (v/v) isotonic Percoll (GE Healthcare) as previously described [37,38]. Enrichment for ring stages following invasion was performed using 5% (w/v) D-sorbitol [38,39].

### Cloning of repair plasmid pT2A-5´UTR-3´-PMV-ΔDHFR

pT2A-5´UTR-3´-PMV-ΔDHFR plasmid is based on vector pT2A-DDI-1cKO (a kind gift of Dr. Edgar Deu, the Francis Crick Institute). The plasmid comprised 5´UTR of *pfpmv* and nucleotides 1-397 of *pfpmv* (Met1 to Lys132), followed by synthetic heterologous *loxP*-containing *sera2* and *sub2* introns (*loxPint*) [24] flanking recodonised *pfpmv* sequence encoding residues Asp133 to Thr590 (GeneArt) fused to a 3xHA epitope tag sequence, a 2A ‘skip’ peptide (2A) [40], and the neomycin resistance gene (*neo*). The second *loxPint* site was immediately followed by another 2A sequence, a *gfp* gene and nucleotides 1333 to 1773 of *pfpmv* which encode Lys445 to Thr590, as a 3´-targeting sequence. Overlapping PCR was used to generate the *loxPint*-2A-*gfp* fragment from the template pT2A-DDI-1-cKO-complement (obtained from Dr. Edgar Deu, the Francis Crick Institute). The resulting PCR product was digested with *Xho*I and ligated into pT2A-DDI-1cKO pre-digested with the same restriction enzymes, yielding plasmid pT2A-DDI-1-cKO-modified GFP (S1 Fig). The 5´-targeting sequence (903 bp) was PCR amplified from 3D7 genomic DNA using Phusion HF DNA polymerase (NEB) with forward primer P3.1-*Bgl*II_5´UTR_F-mod and reverse primer P2-Int_PMV_R. The recodonised fragment (1,602 bp) containing *loxPint*, recondonised *pfpmv*, 3xHA, and 2A was amplified from the plasmid 17ACRILIP_2177297_endoPMV (GeneArt) with primers P4-PMV_int_F and P5-T2A_HA_R. PCR products of 5´-target region and the recodonised fragment were ligated with pT2A-DDI-1-cKO-modified GFP pre-digested with *Bgl*II and *Sal*I using In-fusion^®^ HD cloning kit (Clontech, Mountain View, CA), producing the plasmid pT2A-5´UTR-PMV-cKO. The 3´-target region (441 bp) was amplified from 3D7 genomic DNA using Phusion HF DNA polymerase (NEB) with forward primer P16-EcoRV-PMV-F and reverse primer P17-EcoRV-PMV-R. This fragment was then ligated into pT2A-5´UTR-PMV-cKO predigested with *Eco*RV, yielding the plasmid pT2A-5´UTR-3´PMV-cKO. The *hdhfr* gene was removed from pT2A-5´UTR-3´PMV-cKO by digesting the plasmid with *Bam*HI and *Eco*RI. The plasmid backbone was blunt-end using T4 DNA polymerase (NEB) then religated using T4 DNA ligase (NEB), giving rise to the repair plasmid pT2A-5´UTR-3´-PMV-ΔDHFR.

### Insertion of guide RNA sequences into CRISPR/Cas9 plasmids

Potential guide RNA sequences specifically targeting *pfpmv* were identified using Benchling (https://benchling.com/crispr). Two sets of guide RNA sequences were selected. Two pairs of complementary oligonucleotides (P18-sgPMV-1F and P19-sgPMV-1R; P20-sgPMV-2F, and P21-sgPMV-2R) corresponding to the 19 nucleotides adjacent to the identified PAM sequences were phosphorylated using T4 polynucleotide kinase, annealed and ligated into pL-AJP_004 [41] predigested with *Bbs*I, resulting in the two guide vectors pSgRNA1 and pSgRNA2.

### Generation of *pmv-loxPint* parasites and conditional PMV truncation

The repair plasmid pT2A-5´UTR-3´-PMV-ΔDHFR was linearized with *Sca*I prior to electroporation. Percoll-enriched mature schizonts of *P. falciparum* clone B11 were electroporated with 20 µg of pSgRNA1 or pSgRNA2 and 60 µg of linearized pT2A-5´UTR-3´-PMV-ΔDHFR using anAmaxa P3 primary cell 4D Nucleofector X Kit L (Lonza) as described [22]. Twenty-four hours post-transfection, the electroporated parasites were treated with 2.5 nM WR99210 for 96 h to select for transfectants harbouring pSgRNA plasmids before returning the cultures to medium without drug. Integrant parasites generally reached parasitaemia levels suitable for cryopreservation within 2-5 weeks. Detection of the *pfpmv*-loxPint modified locus was carried out by diagnostic PCR using primer pairs P6-5´UTR_Screen_F and P23-rcPMV-5´integr-R, P6-5´UTR_Screen_F and 198_GFP_start_seq_R, and P24-GFP-3´integr-F and P25-3´UTR-PMV-R. The wild-type *pfpmv* locus was detected by diagnostic PCR using primers P6-5´UTR_Screen_F and P22-PMV-endo-R. Transgenic parasite clones were obtained by limiting dilution cloning by plating a calculated 0.3 parasite per well in flat-bottomed 96-well microplate wells as described [42]. Wells containing single plaques were subsequently expanded into round-bottomed wells. Transgenic parasite clones (PMV-C5 and PMV-F9) were finally checked by diagnostic PCR for integration and modification of the endogenous *pfpmv* gene. Once established, all transgenic clones were maintained in medium without any drug.

Recombination between the *loxPint* sites was induced in tightly synchronised ring-stages of parasite clone PMV-C5 by incubation for 3 h in the presence of 100 nM RAP in 1% (v/v) DMSO; mock treatment was with 1% (v/v) DMSO only. DiCre-mediated excision of the floxed *pfpmv* was detected by PCR analysis of parasite genomic DNA using primers P6-5´UTR_Screen_F and P23-rcPMV-5´integr-R, and P6-5´UTR_Screen_F and Deu198_GFP_start_seq_R. Truncation of *Pf*PMV was evaluated by immunoblot analysis of SDS extracts of mature Percoll-enriched schizonts, probing with the anti-HA antibody 3F10 (Roche), followed by horseradish peroxidase-conjugated secondary antibodies.

### Nucleic acid extraction and polymerase chain reaction

For DNA extraction, total cell pellets were first treated with 0.15% saponin in PBS for 10 min, then washed with PBS before DNA was extracted using a DNeasy^®^ Blood & Tissue Kit (QIAGEN). For diagnostic PCR amplification, GoTaq^®^ (Promega) DNA master mix was used. Amplification of fragments used in construct design was carried out using Phusion^®^ high fidelity DNA polymerase (NEB).

### Indirect immunofluorescence Analysis (IFA) and Western blots

For IFA, thin blood films were prepared from synchronous *P. falciparum* cultures enriched in mature schizonts. The air-dried thin films were fixed in 4% (w/v) paraformaldehyde for 30 min, permeabilised with 0.1% (v/v) Triton X-100 for 10 min, and blocked overnight in 3% (w/v) bovine serum albumin (BSA) in PBS. Slides were probed with rat anti-HA 3F10 (1:100) to detect HA-tagged proteins, human anti-MSP1 monoclonal antibody (mAb) X509 (1:500) to detect MSP1, mAb 89 (1:100) to detect KAHRP, and mAb 2G12 (1:100) to detect HRP2. Primary antibodies were detected using Alexa Fluor 594-conjugated anti-human or anti-mouse secondary antibodies (Life Technologies), and Alexa Fluor 488-conjugated streptavidin (Life Technologies), diluted 1:2000. Slides were stained with 4,6-diamidino-2-phenylindole (DAPI), mounted in Citifluor (Citifluor Ltd., UK). Images were visualised using a Nikon Eclipse Ni microscope with LED-illumination with a 63x Plan Apo λ NA 1.4 objective. Images were taken using an Orca Flash 4 digital camera controlled by Nikon NIS Element AR 4.30.02 software. All images were subsequently analysed using FIJI software.

For Western blots, Percoll-enriched schizonts were pelleted, then resuspended into 10 volumes of PBS. Samples were solubilised into SDS sample buffer, boiled, sonicated and centrifuged. The extracts were subjected to SDS-PAGE under reducing conditions followed by transfer to nitrocellulose membrane. Membranes were probed with rat anti-HA 3F10 (1:1000), human anti-MSP1 mAb X509 (1:1000), anti-GFP (1:1000), mAb 89 (1:1000) or mAb 2G12 (1:1000), followed by horseradish peroxidase-conjugated secondary antibodies. Antigen-antibody interactions were visualised by enhanced chemiluminescence (SuperSignal West Pico chemiluminescent substrate, Pierce).

### Parasite growth assay

Parasitaemia measurement by FACS was as described previously [43]. Briefly, parasites recovered at various time-points were fixed in 8% paraformaldehyde 0.04% glutaraldehyde, pH 7.4 and stained with 2 µM Hoechst 33342 (Invitrogen, Waltham, MA). Parasitaemia was calculated using the FACS BD Fortessa flow cytometer (BD Biosciences). Briefly, cultures to be analysed were initially screened using forward and side scatter parameters and gated for erythrocytes. From this gated population, the proportion of Hoechst-stained cells in 100,000 cells was determined using ultraviolet light with a violet filter (450/50 nm). Samples were analysed using FlowJo software.

## Acknowledgements

The authors are grateful to Sophie Ridewood and Edgar Deu (Francis Crick Institute) for the gifts of pT2A-DDI-1 and pT2A-DDI-1-cKO-complement plasmids and for expert help with flow cytometry analysis.

## Author Contributions

Conceived and designed the experiments: NB, CRC, FH, CWM, MJB. Performed the experiments: NB, CRC, FH. Analysed the data: NB, CRC, MJB. Wrote the paper: NB, CRC, MJB.

